# SARS-CoV-2 spike protein arrested in the closed state induces potent neutralizing responses

**DOI:** 10.1101/2021.01.14.426695

**Authors:** George W. Carnell, Katarzyna A. Ciazynska, David A. Wells, Xiaoli Xiong, Ernest T. Aguinam, Stephen H. McLaughlin, Donna Mallery, Soraya Ebrahimi, Lourdes Ceron-Gutierrez, Leo C. James, Rainer Doffinger, Jonathan L. Heeney, John A. G. Briggs

## Abstract

The majority of SARS-CoV-2 vaccines in use or in advanced clinical development are based on the viral spike protein (S) as their immunogen. S is present on virions as pre-fusion trimers in which the receptor binding domain (RBD) is stochastically open or closed. Neutralizing antibodies have been described that act against both open and closed conformations. The long-term success of vaccination strategies will depend upon inducing antibodies that provide long-lasting broad immunity against evolving, circulating SARS-CoV-2 strains, while avoiding the risk of antibody dependent enhancement as observed with other Coronavirus vaccines. Here we have assessed the results of immunization in a mouse model using an S protein trimer that is arrested in the closed state to prevent exposure of the receptor binding site and therefore interaction with the receptor. We compared this with a range of other modified S protein constructs, including representatives used in current vaccines. We found that all trimeric S proteins induce a long-lived, strongly neutralizing antibody response as well as T-cell responses. Notably, the protein binding properties of sera induced by the closed spike differed from those induced by standard S protein constructs. Closed S proteins induced more potent neutralising responses than expected based on the degree to which they inhibit interactions between the RBD and ACE2. These observations suggest that closed spikes recruit different, but equally potent, virus-inhibiting immune responses than open spikes, and that this is likely to include neutralizing antibodies against conformational epitopes present in the closed conformation. Together with their improved stability and storage properties we suggest that closed spikes may be a valuable component of refined, next-generation vaccines.

## Introduction

The surface of the SARS-CoV-2 virion is studded with spike (S) protein trimers. They are predominantly in a pre-fusion form (Ke et al. 2020; Turoňová et al. 2020) in which the three copies of the receptor binding domain (RBD) are located at the top of the spike, surrounded by the N-terminal domains (NTDs) (Walls et al. 2020; Wrapp et al. 2020). Pre-fusion S trimers are in a stochastic mixture of conformations (Ke et al. 2020), either closed in which all three RBDs lie down at the top of the spike, or open in which one or more of the RBDs protrude from the top of the spike. The Receptor Binding Site (RBS) on the RBD which is responsible for interaction with the receptor ACE2 is largely occluded when the RBD is in the down position (Lan et al. 2020; Shang et al. 2020; Walls et al. 2020; Wrapp et al. 2020). S contains a furin cleavage site, at which it can be separated into S1 and S2 subunits. Cleavage modulates infectivity in a cell type dependent manner (Hoffmann et al. 2020).

After interaction with the receptor, S undergoes a conformational rearrangement leading to exposure of S2, insertion of the fusion peptide (FP) into the membrane of the target cell, and refolding of S2 into the elongated post-fusion form (Cai et al. 2020). This refolding pulls the fusion peptide and transmembrane domain of S together, drawing the target cell and viral membranes together and causing their fusion.

The currently licensed SARS-CoV-2 vaccines in clinical use are designed to deliver the S protein of SARS-CoV-2 as the immunogen (Krammer 2020). The first three licensed vaccines require delivery of either mRNA (Pfizer/BioNtech (Polack et al. 2020), Moderna (Jackson et al. 2020)) or attenuated chimpanzee adenovirus (AstraZeneca/Oxford (Folegatti et al. 2020)) for expression of S protein by target cells in vivo. Vaccine candidates based on inactivated virus, VLP or recombinant protein present the S antigen directly. Since the goal is to generate neutralising antibodies against the virus, a number of the vaccines rely on mutations in S to stabilize the prefusion state to reduce stochastic transition into the postfusion form. Mutations incorporated into current vaccine candidates include the replacement of two residues with a double proline (eg Pfizer/BioNtech (Polack et al. 2020), Moderna (Jackson et al. 2020), Novavax (Keech et al. 2020), Jannsen (Sadoff et al. 2020), as well as mutations in the furin cleavage site for protease resistance (Keech et al. 2020; Sadoff et al. 2020).

Increased stability of S may have advantages in terms of the strength of immune response, particularly in generating antibodies against conformational rather than linear epitopes. Ex vivo, a more stable S may facilitate storage and distribution of protein or VLP vaccines to vaccination sites where cold-chain maintenance is difficult. Krammer and co-workers recently compared the immune response in mice expressing combinations of double proline and cleavage site mutants, finding that both were needed in a recombinant protein vaccine to give complete protection against challenge in the mouse model (Amanat et al. 2020).

Sera from infected individuals contain antibodies against S. Neutralizing antibodies, in particular those against RBD, are being evaluated as antibody therapeutics (Klasse and Moore 2020). Antibodies against RBD include those that directly block the interaction between the RBD and ACE2, some of which are able to bind to RBD in both up and down conformations, and others which only bind the up conformation (Barnes et al. 2020). Some of these antibodies bind between down RBDs in the same trimer, stabilizing the closed form of the spike. Other antibodies against the RBD bind outside of the RBS. Within S there are other epitopes targeted by neutralizing antibodies in the NTD and elsewhere (Gavor et al. 2020; Liu et al. 2020). Polyclonal antibody responses that target divergent epitopes are more robust against escape (Weisblum et al. 2020).

We and others have developed S protein constructs which exclusively adopt closed prefusion conformations (Henderson et al. 2020; McCallum et al. 2020; Xiong et al. 2020), where the RBD should not be accessible to ACE2 binding. Preventing transition to an open state leads to an increase in thermal stability. We speculated that closed spikes would lead to a different polyclonal antibody response when compared to standard S protein constructs, for example by driving the maturation of antibodies that lock the RBD in the closed prefusion conformation

Here we have immunized mice with a range of different S protein constructs that differ in the presence or absence of the double proline mutant, the mutation present at the furin cleavage site, and in whether they are stabilized in the closed prefusion conformation. Comparison of the resulting immune responses revealed that the closed, stabilized spike induces potently neutralising antisera that targets a different pattern of epitopes when compared to more standard stabilized S proteins. These observations suggest that S trimers arrested in the closed state may have value as a component of the next generation of SARS-CoV-2 vaccines. Such vaccines will be designed to induce a broader spectrum of more potent neutralizing antibody responses of different classes with the aim of generating responses effective in protection from evolving circulating variants.

## Results

### Overview of stabilized S protein trimers

We speculated that immunizing with S proteins stabilized in a closed, trimeric conformation would lead to immune responses that differed in strength, and in the range of epitopes targeted, when compared to non-stabilized S trimers, or S trimers that are able to transition into the open form. We therefore set out to compare the immune responses to four constructs: S-GSAS/PP, S-R/PP, S-R, and S-R/x2 (**Fig. 1A**). S-GSAS/PP contains a GSAS sequence preventing cleavage at the furin cleavage site as well as two stabilizing prolines (Wrapp et al. 2020) and provides protection against SARS-CoV-2 infection in mice (Amanat et al. 2020). It is the S antigen of the Adenovirus-expressed Janssen vaccine candidate currently in advanced clinical trials (Mercado et al. 2020) and is the protein component of a vaccine under development by Novavax (Keech et al. 2020). S-R/PP and S-R (Xiong et al. 2020) both have a deletion at the furin cleavage site leaving only a single arginine residue, preventing cleavage. S-R/PP additionally contains the two stabilizing prolines. S-R/x2 contains two cysteine residues at positions 413 and 987 which form a disulphide bond that constrains the trimer in its closed state (**Fig. 1B**) and which results in a dramatic improvement in trimer stability (Xiong et al. 2020).

**Figure 1:**
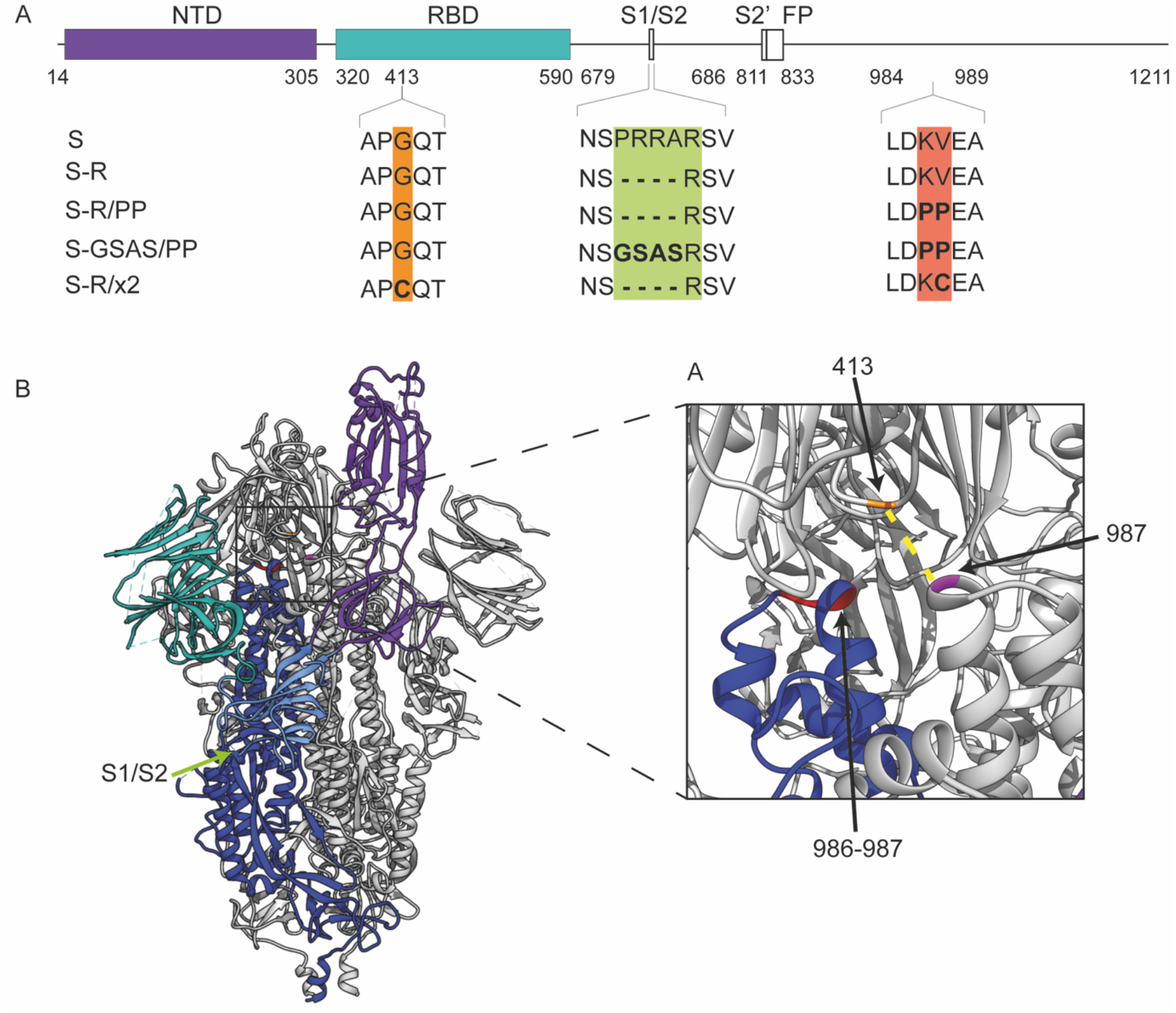
Design of constructs. **A.** Overview of constructs used in the study indicating the positions where residues have been mutated. **B.** An overview of the trimeric spike structure indicating the positions of the mutated residues. The insertion of cysteine residues at positions 413 and 987 leads to the formation of a disulphide bond (dotted yellow line in inset) that arrest S in the closed prefusion form.

S trimers were expressed as described previously (Xiong et al. 2020). Proteins were purified by metal affinity chromatography as described previously (Xiong et al. 2020) and quality controlled by negative stain electron microscopy.

### S-R/x2 is arrested in a closed state, leading to reduced ACE2 binding and reduced infectivity

To assess the impact of the disulphide bond in S-R/x2 on S-mediated viral entry, we infected HEK293T-hACE2 cells with S-pseudotyped, replication-deficient lentiviruses. Pseudovirions bearing S-R/x2 showed minimal infection compared to pseudovirions bearing wild-type S or S-R (**Fig. 2A**). Deletion of the C-terminal 19 amino acids from S substantially increased infection by pseudovirions bearing wild-type S, consistent with previous data (Ujike et al. 2016), and increased infectivity of S-R pseudovirions. However, pseudoviruses with C-terminally deleted S-R/x2 gave minimal infection (**Fig. 2A**). These data demonstrate that stabilization of S in the closed state via the x2 disulphide renders the virus largely non-infectious, consistent with the RBS in the RBD being inaccessible for ACE2 binding in the closed state.

**Figure 2.**
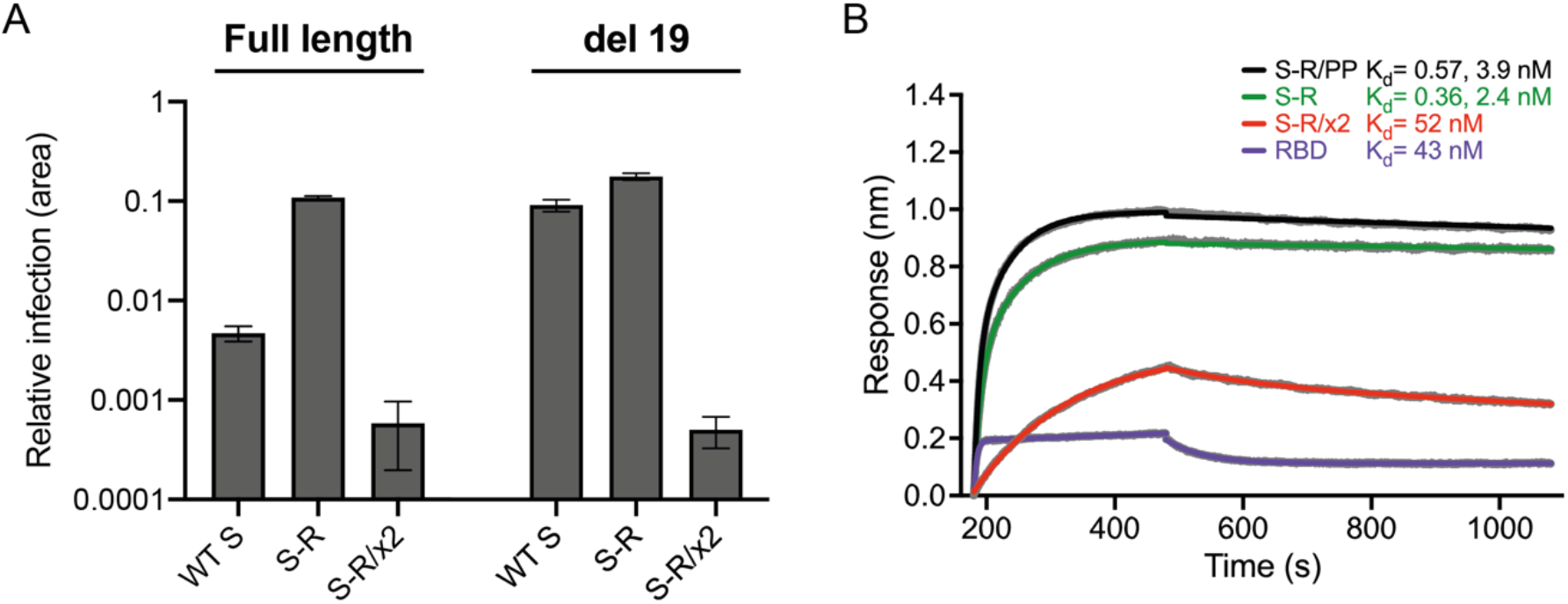
Characterization of fusogenicity and ACE2 binding. **A.** Infection of HEK293T-hACE2 cells with SARS-CoV-2 S-pseudotyped HIV virions carrying a GFP reporter gene. The relative area of infected cells was quantified by GFP fluorescence 48 h post-infection using an Incucyte S3 live cell imager. Viruses were produced using either full-length S constructs or a C-terminal deletion of 19 amino acids (del 19) to increase infectivity. Virions pseudotyped with wild-type (WT) S are compared to those pseudotyped with S-R and S-R/x2. **B**. Biolayer interferometry sensograms for binding kinetics of ACE2 to 600 nM RBD (magenta), and 1000 nM of S-R/PP (black), S-R (green) and S-R/x2 (red). Data are shown in grey with fits to the data in their respective coloured lines. The dissociation constants (*K_d_*) shown were calculated from the fitted association (*k_on_*) and dissociation (*k_off_*) constants for the interactions summarised in Table 1, where *K_d_* = *k_off_/k_on_*, from concentration dilution series (**Fig. S2**).

We next investigated how stabilizing S in the closed state affected the kinetics of the interaction with ACE2 using biolayer interferometry (**Fig. 2B, Table 1, Fig. S1**). Free RBD associated and dissociated relatively rapidly with a binding constant of 43 nM consistent with previous observations (Wrapp et al. 2020). S-R and S-R/PP displayed more complex kinetics, with two association phases: one in the same order as free RBD, and one slower, suggesting either different conformations of S or a secondary rate-limiting step such as a conformation change. The dissociation rate was 3 orders of magnitude slower than free RBD leading to a half-life of the complex of between 2.5 to 5 h and resulting in apparent binding constants in the low nM and high pM range (**Table 1**). The slow dissociation rate is likely due to avidity from the three RBDs on the S trimer. The interaction kinetics of S-R/x2 with ACE2 were significantly altered with a single association rate two orders of magnitude slower than free RBD and the other S proteins, and a dissociation rate an order of magnitude faster than the other S proteins. The change in interaction kinetics results in formation of fewer S – ACE2 complexes (approximately half the amplitude observed for the other S proteins), and the half-life of complexes is reduced to approximately 35 mins. Hence the binding affinity of S-R/x2 to ACE2 is between 10 and 100 times weaker than the other S proteins, consistent with the ACE2 binding site being largely hidden in S-R/x2.

**Table 1:** Summary of fitted association (*k_on_*) and dissociation (*k_off_*) constants for the ACE2 interactions. K^*kin*^_*d*_, the kinetic dissociation constant, = *k_off_/k_on_* from concentration dilution series (see Fig. S2).

| Ligand | Analyte | $k_{on} (M^{-1}s^{-1})$ | $k_{off} (s^{-1})$ | $K^{kin}_d (nM)$ |
| --- | --- | --- | --- | --- |
| ACE2-Fc | RBD | $3.8 \times 10^5$ | 0.016 | 43 |
| | S-R/x2 | $6.3 \times 10^3$ | $3.3 \times 10^{-4}$ | 52 |
| | S-R | $1.0 \times 10^5$ (fast) | $3.6 \times 10^{-5}$ | 0.36 |
| | | $1.5 \times 10^4$ (slow) | | 2.4 |
| | S-R/PP | $1.3 \times 10^5$ (fast) | $7.4 \times 10^{-5}$ | 0.57 |
| | | $1.9 \times 10^4$ (slow) | | 3.9 |

### Immunization

S trimers were thawed 30 minutes prior to immunization and were mixed with the adjuvant Monophosphoryl lipid A (MPLA) in phosphate buffered saline. Mice were immunized with 10 μg of protein and 10 μg MPLA then boosted 4 weeks after the first immunization with 10 μg protein and 2 μg MPLA, according to the protocol illustrated in **Fig. 3A**. Mice were bled 2, 4, 6, 8 and 21 weeks post immunization (**Fig. 3A**) and culled 25 weeks post immunization by cardiac puncture under terminal anaesthesia. Adjuvant-mixed trimers were imaged by negative stain EM, and in all cases large numbers of trimeric S proteins were visible, validating that S trimers were intact in the presence of adjuvant at the point of immunization (**Fig. 3B**).

**Figure 3.**
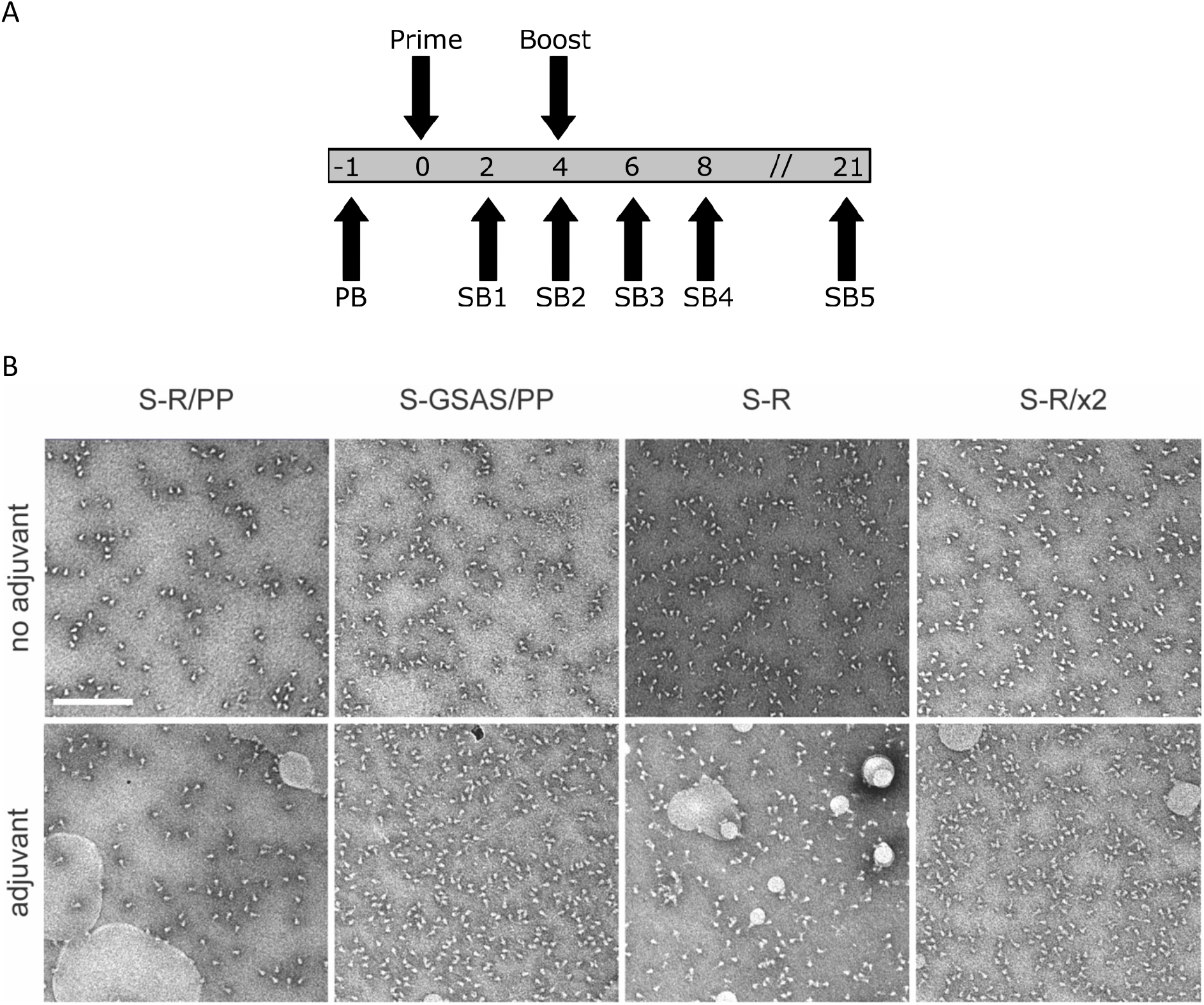
Immunization strategy and proteins for immunization. **A.** Overview of immunization strategy. 8-10 week old BALB/c females were immunised twice with a four week interval. Mice were bled 1 week prior to immunisation (PB), and serial bleeds (SB) were taken at two weekly intervals after the first immunisation, and again at 21 weeks, after which animals were sacrificed and spleens taken for T-cell assays. **B.** Oligomeric state of S proteins before and after addition of adjuvant assessed by negative stain electron microscopy. In all cases the proteins are predominantly in the prefusion, trimeric state. Scale bar 200 nm.

### Trimeric S constructs induce neutralizing sera in mice

We assessed the degree to which sera were able to neutralize S-mediated infectivity in a lentiviral pseudovirus assay. The majority of mice immunized with S variants had neutralizing sera two weeks post-immunization, and by two weeks post-boost the sera were potently neutralizing, with the geometric mean of 50% pseudovirus neutralisation titre for each variant in the range of 7043-12122 (**Fig. 4A, Fig. S2**). Sera remained potently neutralizing 21 weeks post immunization. There were no statistically significant differences in the neutralizing titre between the different trimeric S variants assessed at any time point post-immunization, however average neutralising antibody titres for S-R/x2 and S-GSAS/PP were higher than those for S-R and S-R/PP in the functional assays.

**Figure 4.**
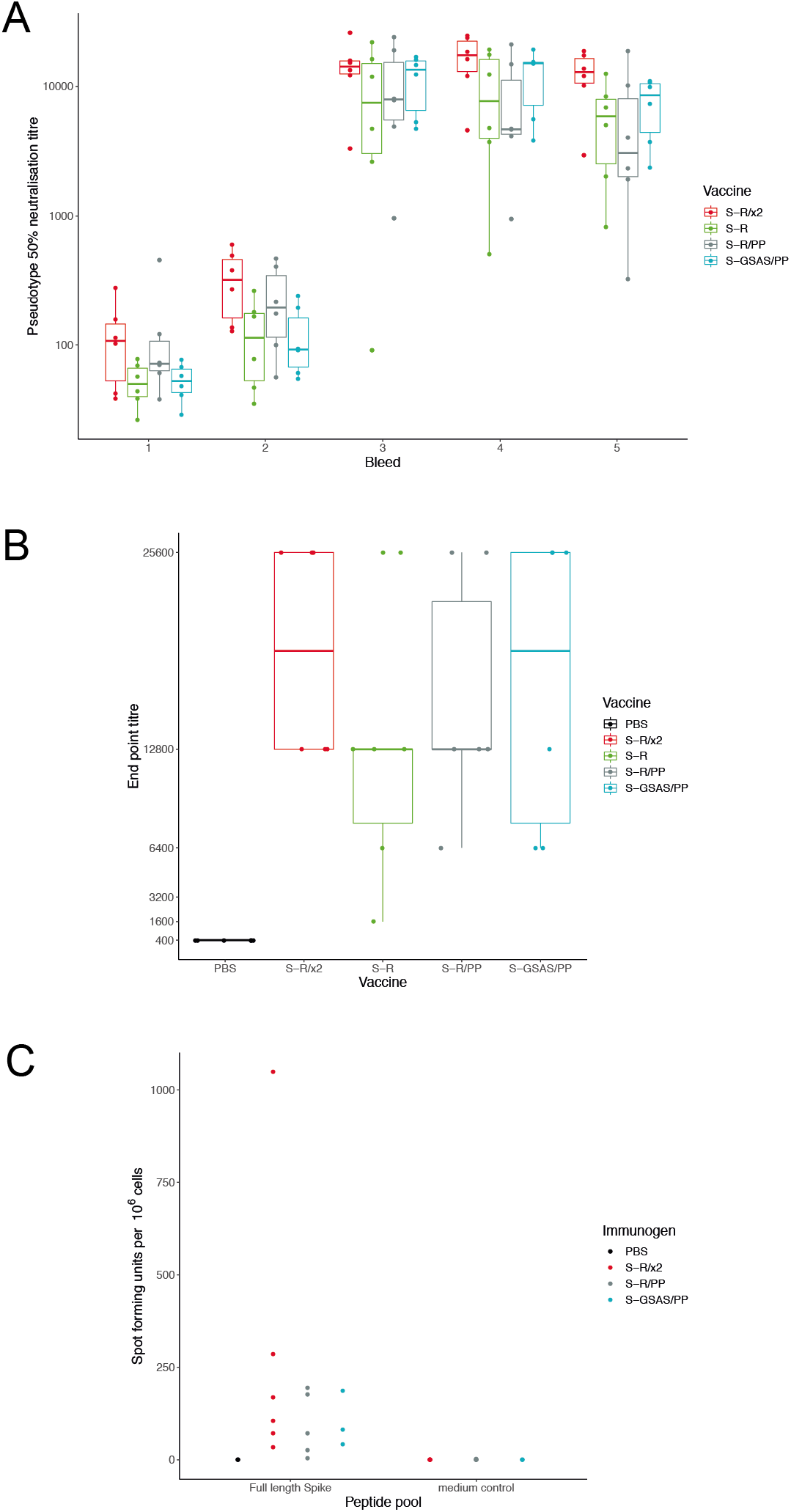
Neutralisation by sera and T-cell response. **A.** SARS-CoV-2 pseudovirus 50% neutralisation titre for individual mice at five bleed points post immunisation with one of four S protein variants. Bleed timepoints are 2, 4, 6, 8, and 21 weeks post prime. **B.** box whisker plots for endpoint antibody titres representing the serum dilution point at which the cytopathic effect in Vero cells caused by SARS-CoV-2 BetaCoV/Australia/VIC01/2020 infection was no longer inhibited, as compared to controls. Endpoint titre means ranged between 12800 and 25600 for vaccine sera. **C.** Comparison of persisting T-cell responses 21 weeks post vaccination with stabilised S constructs. T-cell responses were generated by all constructs with some heterogeneity: spot forming units range from 0-88 for S-R/PP. 13-79 for S-GSAS/PP and 31-424 for S-R/x2.

To assess the ability of the sera to neutralize infectious SARS-CoV-2 virus, we performed virus neutralization assays against SARS-CoV-2 BetaCoV/Australia/VIC01/2020 (Caly et al. 2020) and measured endpoint titres for sera collected at bleed 4 (**Fig. 4B**). Cytopathic effect-based endpoint neutralising antibody titres ranged from 1600 to 25,600 with the S-R/PP average endpoint titre at 16,000, S-GSAS/PP at 17,066, S-R at 12,000 and S-R/x2 at 19,200. These are substantially higher than endpoint titres from a standard high antibody titre convalescent human sera control (NIBSC 20/130, see methods) which ranged from 1600 to 6400 with an average of 3466.

### Trimeric S constructs induce T-cell responses

To assess whether immunization led to the development of S-protein-specific T cells, we stimulated splenocytes from vaccinated mice with peptide pools covering the complete S protein and analysed the response using an IFN-g ELIspot assay (**Fig. 4C, Fig. S3**). We analysed T cells from sets of 3 (or 6) mice immunized with S-R/GSAS, S-R/PP and S-R/x2. Though low to moderate, T-cell responses to these different forms of S proteins adjuvanted in MPLA, were consistently detected in all immunized mice.

### Trimeric S constructs all induce antibodies that block the ACE2-RBD interaction in vitro

We next explored whether the S protein constructs induced antibodies that are able to directly block the RBD-ACE2 interaction using a surrogate-neutralization assay (Genscript, (Tan et al. 2020)). Sera from bleed 3 were incubated with Horse radish peroxidase-coupled RBD, which was then bound to ACE2-coated wells before enzymatic read out of the degree of binding. All S-immunized sera inhibited RBD-ACE2 interactions, indicating that both open and closed S protein constructs can induce antibodies that directly block the interaction between ACE2 and the RBS on the RBD (**Fig. 5A**). These observations are consistent with the exposure of part of the RBS in the closed conformation adopted by S-R/x2 trimer. We compared the inhibition of RBD-ACE2 interactions with neutralization in the pseudovirus based assay (**Fig. 5B**). These variables are well correlated, and this comparison revealed that sera from S-R/x2-immunized mice neutralized virus entry better than would be expected based on their inhibition of the RBD-ACE2 interaction. This observation suggests that the potent neutralization by sera from S-R/x2 immunized mice involves antibodies that prevent S-mediated fusion without directly blocking interaction between the RBS and the receptor. Such antibodies might include those that bind to the RBD and prevent S opening by conformationally stabilizing the closed state, or that bind to other epitopes on S and thereby inhibit receptor interactions or fusion.

**Figure 5.**
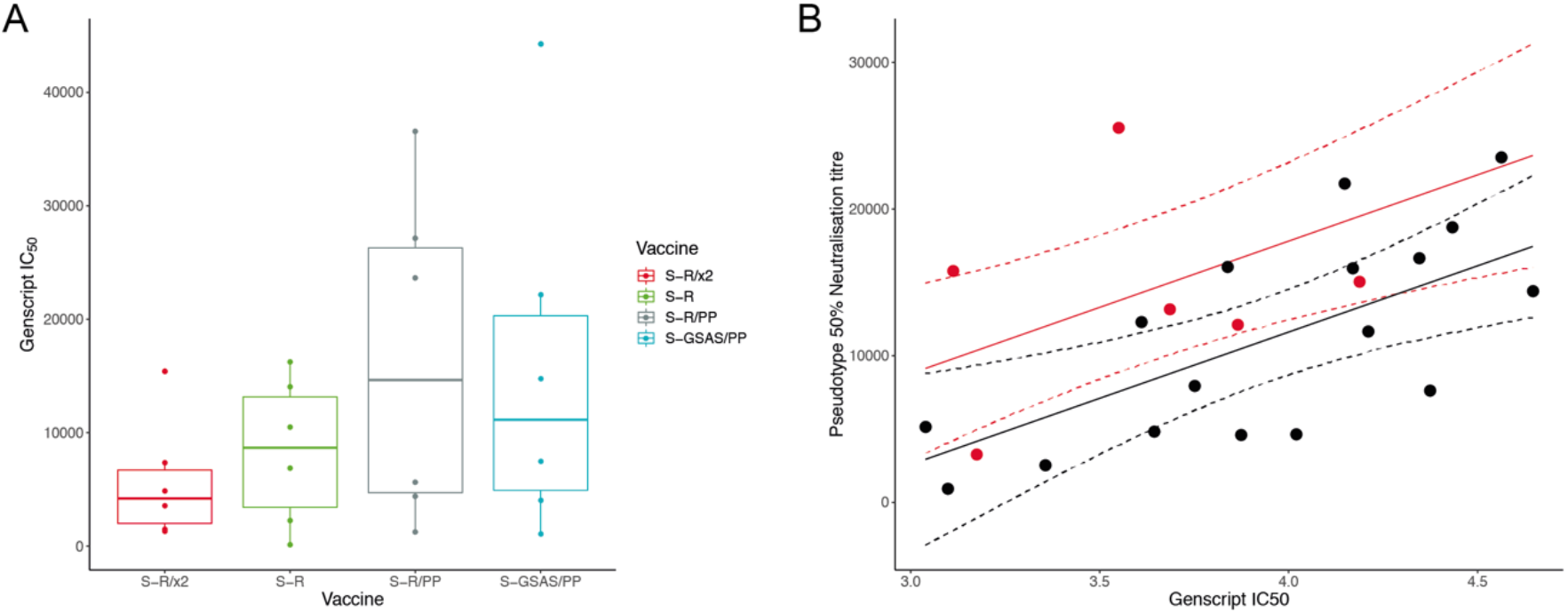
Inhibition by sera of RBD-ACE2 interaction. **A.** Boxplot showing IC_50_ values for the Genscript assay measuring inhibition of RBD-ACE2 interactions by individual mice sera. **B.** Sera from S-R/x2 immunised mice were significantly more potent neutralizers of SARS-CoV-2 pseudovirus than expected based on their direct inhibition of RBD-ACE2 interaction. Predicted neutralisation shown in solid line, 95% confidence intervals shown with dashed lines. S-R/x2 sera and predictions shown in red, all other sera shown in black.

### Sera contain antibodies against linear epitopes from surface-exposed positions on S

For a set of six mice, three immunized with S-R/PP and three immunized with S-R/x2, we performed peptide array analysis using overlapping 15mer peptides covering the complete SARS-CoV-2 S to identify linear epitopes bound by antibodies in the sera (**Fig. 6A**). The pattern of linear epitopes varied between sera. They represented almost exclusively surface accessible residues and were predominantly accessible loops or strands (**Fig. 6B**). Among the most strongly bound epitopes were those in the vicinity of heptad repeat 2 (HR2) and the fusion peptide (FP) which are also widespread in sera from infected humans (Garrett et al. 2020; Poh et al. 2020; Shrock et al. 2020). We identified only one strongly binding internal linear epitope, which is centered around residue 991 in a helix, and which bound sera from one of the S-R/PP immunized mice. This epitope is unlikely to be accessible in the folded prefusion spike, suggesting the presence of some unfolded S-R/PP after immunization.

**Figure 6.**
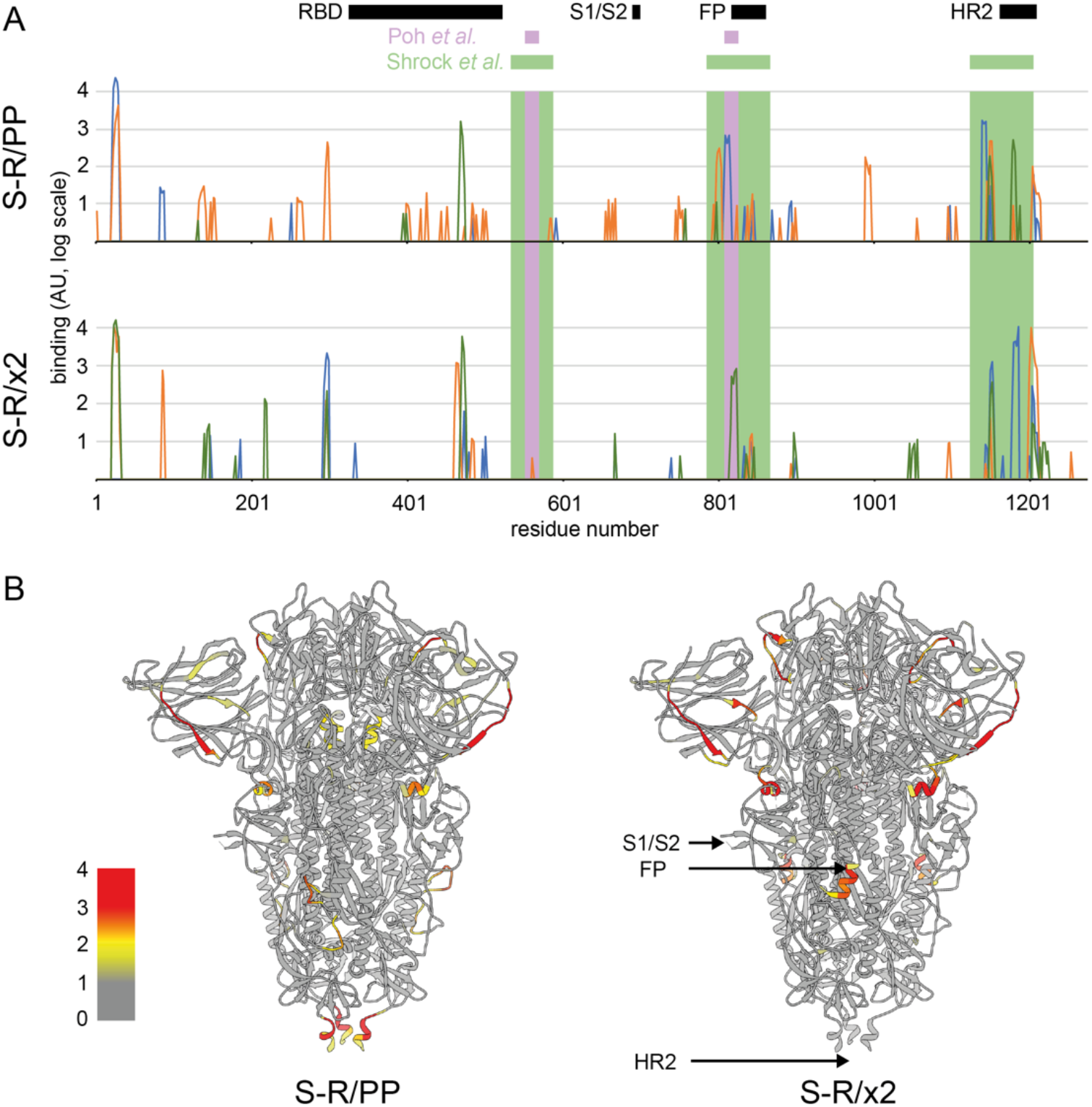
Quantification of binding of sets of three sera immunized with S-R/PP or S/R/x2 to an array of overlapping 15mer linear peptides covering S. **A.** the binding strengths of different sera are shown as blue, green and orange lines. X axis is the residue number at the centre of the peptide, Y axis is the intensity of binding shown on a log scale. Different sera bind different patterns of peptides. The positions of the most common linear epitopes identified in human sera by (Shrock et al. 2020) are shown as pale green rectangles; the positions of two linear peptides that induce a neutralizing antibody response, as identified by (Poh et al. 2020), are shown as purple rectangle. Structural features are indicated: RBD, S1/S2 cleavage site, fusion peptide (RP) and the heptad repeat 2 region (HR2). **B.** The positions of the bound epitopes are coloured on the structure of closed, trimeric S, showing the maximum binding value of the three sera for S-R/PP or S-R/x2. The bound epitopes are almost entirely surface exposed.

### Trimeric S constructs all induce antibodies that bind S constructs and RBD, but S-R/x2 immunization leads to different antibodies

Human sera from infected patients contain antibodies against both tertiary and quaternary structural epitopes as well as the linear epitopes that are represented in peptide arrays. To gain further insights into the sera, we used a multiplexed, particle-based flow cytometry method (Luminex) to measure binding of sera to a range of different S protein antigens: S-GSAS/PP, S-R/PP, S-R, S-R/x2 and RBD (**Fig. 7A**). Sera bound to all of these antigens. There were no significant differences between binding measured to S-GSAS/PP, S-R/PP, S-R, and RBD for sera from mice that had been immunized with different S proteins. Sera from mice that had been immunized with S-R/x2 did, however, bind more strongly to S-R/x2 antigen than sera from other mice. Within the multiplexed assay, we additionally included S-R and S-R/x2 that had been pre-incubated at 37°C for 36 h or 60°C for 30 minutes to induce partial disassembly (see below)

**Figure 7.**
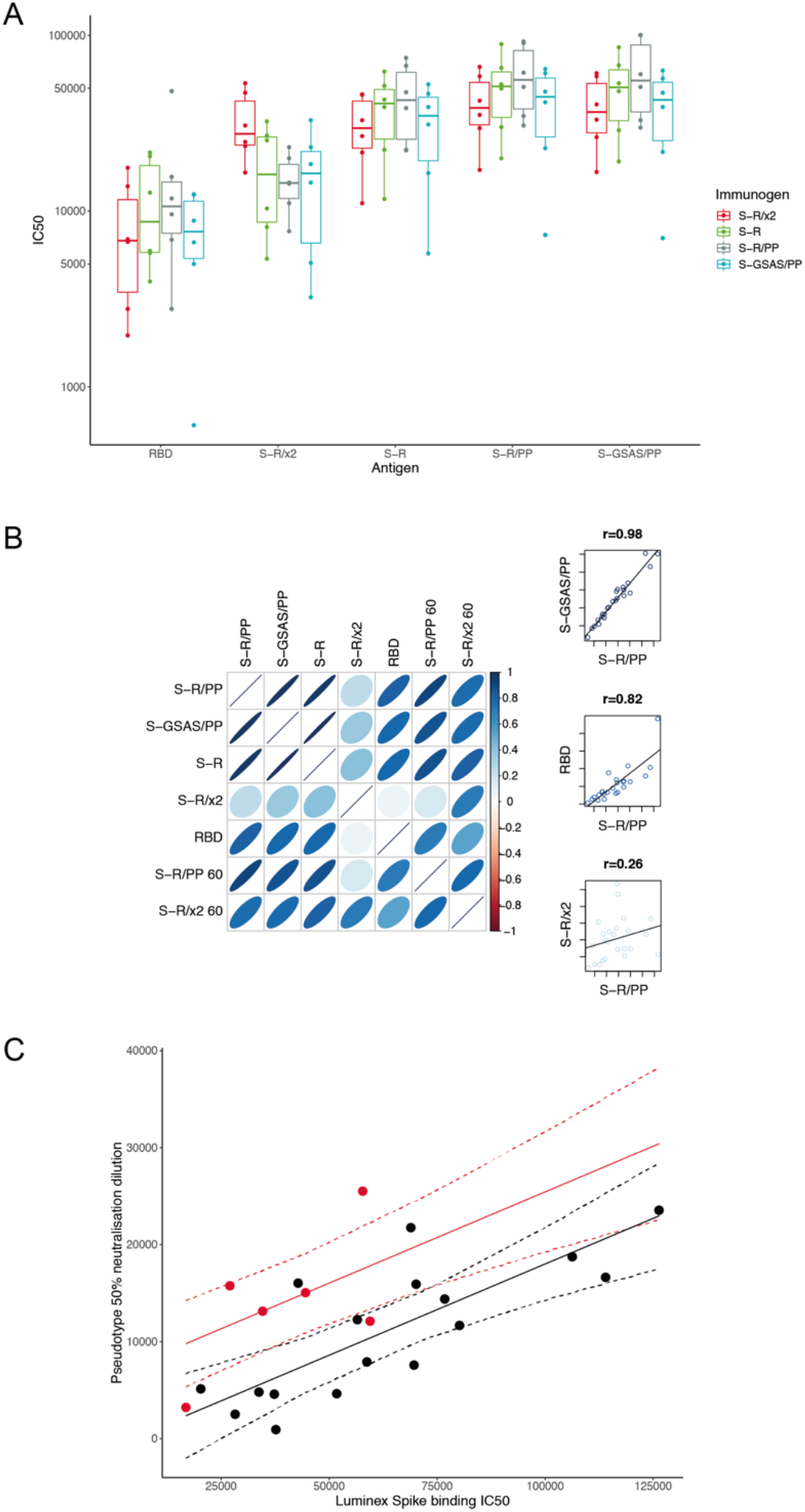
Binding of sera to S protein antigens. **A.** IC_50_ values of sera from individual mice immunised with one of four trimeric S constructs binding to five separate antigens. The antigens are SARS-CoV-2 RBD and the four trimeric S constructs. Binding of sera from mice immunized with different S proteins, to different S antigens, were measured using the Luminex assay. All trimeric S constructs induce antibodies that bind all constructs and RBD. Sera from mice immunized with S-R/x2 bound the S-R/x2 antigen more than sera from other mice. B. Visualisation of correlation in IC_50_ values between different antigens. Narrower, darker ellipses indicate stronger correlations. The three scatter plots on the right hand side are illustrative examples showing the raw relationship for high, low, and intermediate correlations. There is generally a good correlation between strength of individual sera binding to different S antigens. This correlation is less strong for RBD, and very different for S-R/x2, suggesting that S-R/x2 is displaying different epitopes in this assay than the other S proteins. Heating S-R/x2 leads to it displaying epitopes that are more similar to those of the other S proteins. C. Sera from S-R/x2 immunised mice were significantly more potent neutralizers of SARS-CoV-2 pseudovirus than expected based on their Spike binding IC_50_. Predicted neutralisation shown in solid line, 95% confidence intervals shown with dashed lines. S-R/x2 sera and predictions shown in red, all other sera shown in black.

As expected there is very strong correlation between the binding of individual sera to S-GSAS/PP, S-R/PP and S-R (p>=0.97), reflecting the structural similarities between these antigens (**Fig. 7B**). There is also substantial correlation between binding to RBD and to these S variants (p between 0.75 and 0.8), suggesting that the RBD contains immunodominant epitopes, consistent with observations using human sera (Piccoli et al. 2020; Yu et al. 2020; Zhu et al. 2020). In contrast, there is poor correlation (p<0.5) between the binding of individual sera to S-R/x2 and to the other S variants or RBD. This suggests that different epitopes are accessible in the S-R/x2 antigen. S-R/x2 that was previously incubated at 60°C for 30 minutes, which we have previously shown induces partial disassembly (Xiong et al. 2020), binds sera to a degree that correlates better with binding to other S-variants (p between 0.7 and 0.8) than unheated S-R/x2. This implies that after heating, the S-R/x2 antigen exposes additional epitopes and loses other epitopes in a manner that makes it more similar to the other S antigens. We speculate that heating of S-R/x2 leads to exposure of linear epitopes, exposure of epitopes in the RBD that are hidden in the closed conformation, as well as the loss of structural epitopes that are otherwise best preserved in the more stable S-R/x2 protein.

Sera from S-R/x2 immunized mice show increased binding to S-R/x2 antigen, but do not show increased binding to the other S protein antigens. We compared the binding to S-R/PP (as a representative standard S) to pseudovirus neutralization. Sera from mice immunized with S-R/x2 neutralize more strongly than would be predicted based on their binding to S-R/PP (**Fig. 7C**). Considered together with the above observations, these data suggest that immunization with S-R/x2, compared to immunization with other S constructs, leads to a higher fraction of neutralizing antibodies against epitopes which are prevalent in the stable S-R/x2 trimer. We consider it most likely that these are conformational epitopes specific to the closed state. This interpretation explains the observations that S-R/x2 induced sera have increased binding to S-R/x2 antigen, and that they neutralize better than expected given their binding to S.

## Discussion

In summary, immunization of mice with trimeric S protein constructs induces very strongly neutralizing sera, and the sera remain neutralizing 5 months post immunization with neutralising antibody titres maintained at high levels. Immunization also induces a T-cell response. All of the sera contain antibodies that bind S and RBD, and which block the ACE2-RBD interaction, including sera induced by the stabilized, closed S-R/x2 construct.

The poor correlation between binding of sera to S-R/x2 and to other trimeric S constructs, indicates that S-R/x2 displays a different pattern of epitopes when used as an antigen in Luminex assays. Partly this will be due to conformational differences, with S-R/x2 being exclusively in the closed conformation. The observation that heating of S-R/x2 makes it more similar to the other S protein constructs suggests that the S-R/x2 has a more stable quaternary structure than the other antigens, and this more stable structure may also contribute to the different pattern of epitopes.

Our observation that sera induced by the more stable S-R/x2 show significantly better binding to the closed form of the S antigen than sera induced by open or intermediate forms of S trimeric spikes indicates that S-R/x2 raises a different pattern of antibodies compared to other trimeric spikes when used as an immunogen. We suggest that this is for two reasons. Firstly, because of conformational differences: S-R/x2 is exclusively in the closed state, while the other S protein constructs also adopt open states. S-R/x2 therefore does not display epitopes hidden in the closed state, particularly some in the RBD. Secondly, because S-R/x2 is more stable and therefore is likely to remain in a folded state for longer post immunization – for this reason it may raise more conformational antibodies than S-R. Sera induced by S-R/x2 neutralize better than expected based on their binding to other trimeric spike constructs, suggesting that they indeed contain increased neutralizing antibodies specific to the closed conformation. We therefore conclude that S-R/x2 induces more neutralizing conformational antibodies against the closed state than the other trimeric immunogens.

Here we have studied the immune response in naïve mice that do not express human ACE2. S-R/x2 binds ACE2 with markedly reduced affinity than the other S protein constructs we have assessed. This may lead to enhanced differences when used to immunize species with homologous ACE2 receptors.

On an operational level, the increased thermal stability of S-R/x2 may have important advantages for widepread global distribution, given that cold-chain maintenance is logistically challenging. While we have demonstrated the unique properties of S-R/x2 as a protein immunogen, the S-R/x2 sequence or derivatives thereof can readibly be delivered as DNA or RNA as part of next generation vaccine platforms. The increased thermostability of S-R/x2 may still be useful in genetically encoded vaccines due to persistence of the folded state when the gene is expressed in vivo.

New SARS-CoV-2 variants are emerging as the virus evolves increased transmissibility. These include changes in the RBS which increase affinity with human ACE2 and may represent species adaptation. New variants also result from antibody selection pressure. Escape from single neutralising SARS-CoV-2 mAbs to S has been demonstrated *in vitro* (Baum et al. 2020), and neutralisation escape has also been observed in an immunocompromised individual treated with convalescent plasma (Kemp et al. 2020). While dominant neutralisation escape mutants have not yet been observed in the human population, virus escape from current S based vaccines remains a concern as the first generation vaccines are being rolled out. Furthermore, antigen designs to avoid the possibility of inflammatory triggers such as antibody dependent enhancement commonly observed amongst coronaviruses remain important considerations in vaccinating human populations at risk from severe COVID-19 disease. In the context of immunisation of humans previously infected with SARS-CoV-2, or indeed with prior immunization, evaluation of next generation S antigen candidates to recruit additional beneficial neutralising responses will be crucial. In the light of all these concerns, there is a need to expore new immunogens for SARS-CoV-2 vaccination.

The S proteins expressed by the Pfizer/BioNtech and Moderna mRNA vaccines, the AstraZeneca ChAd vaccine, and others, are all based on early S proteins designs that are able to adopt both open and closed conformations. The S-R/x2 closed spike induces a strongly neutralizing response against a different pattern of epitopes than these constructs. This response may be less sensitive to recent changes in the RBS at sites which are largely occluded in the closed form. Engineered closed S constructs may therefore be valuable immunogens for inducing neutralising antibodies against different epitopes. Having S scaffolds such as S-R/x2 that induce polyclonal responses against a different pattern of epitopes allows for flexibility of vaccination regimens at both the individual and population level.

## Acknowledgements

The research reagent NIBSC 20/130 was obtained from the National Institute for Biological Standards and Control, UK. GC and JLH thank Dr Nigel Temperton (University of Kent, UK) for the pCAGGS_TMPRSS2, pCAGGS_hACE-2, p8.91 and pCSFLW plasmids, and Dr Edward Wright (University of Sussex, UK) for the HEK293T/17 cell line.

This study was supported by funding from the European Research Council (ERC) under the European Union’s Horizon 2020 research and innovation programme (ERC-CoG-648432 MEMBRANEFUSION to JAGB), the Medical Research Council as part of United Kingdom Research and Innovation (MC_U105181010 to LCJ; MC_UP_1201/16 to JAGB). Vaccine studies in animals were supported by the UKRI DIOS-Coronavirus vaccine grant (72845 to JLH); and the Humoral Immune Correlates for COVID-19 (HICC, UKRI/NIHR ref COV0170 to JLH), University of Cambridge.

## Materials and Methods

### Protein expression and purification

All S-protein constructs have been previously reported (Xiong et al. 2020). Proteins were expressed and purified exactly as described in Xiong et al 2020.

### Lentiviral pseudotyping to assess infectivity of disulphide-stabilised variant

Vectors to produce pseudovirions were pCRV-1 (encoding HIV-1 Gag Pol) (Naldini et al. 1996), CSGW (encoding GFP) (Zennou et al. 2004) and pCAGGS-S (encoding spike or associated mutant). pCAGGS-S was generated by cloning codon optimised S into a pCAGGS empty backbone using EcoRI and NheI restriction sites. pCAGGS-S Δc19 was generated by deleting the C-terminal 19 amino acids, which have previously been shown to contain an endoplasmic reticulum (ER)-retention signal (Ujike et al. 2016). Replication deficient SARS CoV-2 pseudotyped HIV-1 virions were produced in HEK293T wild-type (293T-wt) cells by transfection with pCAGGS-S, pCRV and CSGW as described previously (Price et al. 2014). Viral supernatants were filtered through a 0.45 μm syringe filter at 48 h post-transfection and pelleted through a 20% sucrose cushion for 2 h at 28000 rpm. Pelleted virions were resuspended in OptiMem.

HEK293T-hACE2 cells were plated into 96 well plates at a density of 7.5×10^3^ cells per well and allowed to attach overnight. Lentiviral pseudotype stocks were titrated in triplicate by addition of virus onto cells. Infection was measured by GFP expression using an Incucyte Live cell imaging system (Sartorius). Infection was enumerated as GFP positive cell area.

### Biolayer interferometry

The pCD5-hACE2-Fc expression vector was a gift from Berend Jan lab. The ACE2-Fc fusion protein was expressed in Expi293 cell and purified from cell culture media using a protein A column followed by size-exclusion chromatography. Ace2-Fc at 11 μg/mL was immobilised onto Protein A BioFsensors to a level of approximately 0.8 nm in a Octet RED384 (FortéBio). The sensor tips were dipped into 1:3 serial dilutions of RBD or S proteins from initial concentrations of 600 and 1000 nM respectively, for 5 mins to observe association followed by transfer to wells containing only assay buffer (10 mM HEPES pH 7.5, 150 mM NaCl, 3 mM EDTA, 0.05% Tween 20 and 1 mg/mL bovine serum album) to follow dissociation for 10 min. The assay was run at 30 °C. The sensor tips were regenerated with 10 mM glycine pH 1.5 and the assay preformed again without loading of ACE2-Fc to monitor non-specific interactions. The data were double-referenced, subtracting a buffer reference and the parallel assay without ACE2-Fc. Resultant reference-subtracted data were fitted to single or double phase association and dissociation kinetics to determine *k*_on_, *k*_off_ and *K*_d_^(kin)^ (the binding constant determined from the ratio of the individual rate constants) using Prism 8.4.3 (GraphPad Software).

### Negative stain EM

To characterise protein samples with adjuvant, 10 μg of protein was mixed with 10 μg monophosphoryl lipid A in 50 μl final volume of PBS. 3 μl of sample diluted to 0.05 mg/ml in water was applied to glow discharged (45s 30 mA) CF200-Cu carbon film grids and absorbed for 30 seconds. The grids were side-blotted and washed three times with water, then stained with Nano-W stain (Nanoprobes) followed by blotting immediately. The grids were air-dried and imaged using a Tecnai T12 microscope operated at 120 kV.

### Animal work

8-10 week old BALB/c females (Charles River) were immunized subcutaneously with 10 μg of purified protein and 10 μg of MPLA in a total volume of 50 μl PBS. Mice were boosted after a four-week interval with 10 μg of purified protein and 2 μg of MPLA. Serial bleeds were taken via the saphenous vein at two-weekly intervals at D0, D14, D28, D42, D56, and D147. Spleens were removed from culled mice for ELISpot analysis.

### Cells and viruses for neutralisation assays

Vero (ATCC CCL 81) and HEK293T/17 (ATCC CRL-11268) cells were cultured in Dulbecco’s modified Eagle Medium, supplemented with penicillin (100 U/ml) and streptomycin (100 μg/ml) and 10% foetal bovine serum. SARS-CoV-2 (BetaCoV/Australia/VIC01/2020) was obtained from Porton Down, Public Health England, and propagated in Vero cells under BSL-3 conditions. Lentiviral pseudotypes were produced by transient transfection of HEK293T/17 cells with packaging plasmids p8.91 (Naldini et al. 1996; Zufferey et al. 1997) and pCSFLW (Zufferey et al. 1998) and a SARS-CoV-2 spike expression plasmid using the Fugene-HD transfection reagent. Supernatants were taken after 48 h, filtered at 0.45 μm and titrated on HEK293T/17 cells transiently expressing human ACE-2 and TMPRSS2.

### Pseudotype-based microneutralisation assay

Pseudotype-based microneutralisation assay was performed as described previously (Carnell et al. 2017). Briefly, serial dilutions of serum were incubated with SARS-CoV-2 spike bearing lentiviral pseudotypes for 1 h at 37°C, 5% CO_2_ in 96-well white cell culture plates. 1.5×10^4^ HEK293T/17 transiently expressing human ACE-2 and TMPRSS2 were then added per well and plates incubated for 48 h at 37°C, 5% CO_2_ in a humidified incubator. Bright-Glo (Promega) was then added to each well and luminescence read after a five-minute incubation period. Experimental data points were normalised to 100% and 0% neutralisation controls and non-linear regression analysis performed to produce neutralisation curves and IC_50_ values.

### Virus neutralisation assay

Vero cells were plated in 96-well clear cell culture plates to reach confluence on the day of infection. Serial dilutions of serum in PBS were incubated with 500 pfu/well of BetaCoV/Australia/VIC01/2020 in PBS for 1 h at 37°C, 5% CO_2_ in a humidified incubator before being transferred to 96-well Vero cell monolayers. Plates were then incubated at room temperature on a rocking platform. After 90 minutes 100 μl of 2x DMEM (2% final FBS) was added and plates incubated at 37°C, 5% CO_2_ in a humidified incubator. After 48h, media was removed and cells fixed with 10% formalin for 30 minutes at room temperature. Cells were then stained with 0.25% crystal violet solution and endpoint dilutions measured.

### Murine Interferon-*γ* ELISpot Assay

Using the flat edge of a 5 mL sterile syringe stopper head, the spleens of freshly sacrificed mice were passed through a 70 μm cell strainer fitted onto a 50 ml falcon tube. Smashed spleen tissue was then washed down through the cell strainer using PBS warmed to room temperature and the resulting cell suspension centrifuged at 330 xg for 5 minutes. To remove red blood cells, the cell pellet was resuspended in 2 ml PBS, overlaid on an equal volume of Histopaque^®^ 1083 (Sigma-Aldrich, 10831-100M) and centrifuged at 400 xg for 30 minutes without brake. The layer of white cells was carefully extracted and washed at 330 xg for 5 minutes in RPMI-1640 (Gibco, 21875-034) prewarmed to 37°C. The cell pellet was resuspended in prewarmed (37°C) complete CTL-TestTM medium and live cells were counted. Cell concentration was adjusted to 4 x 10^6^ cell/ml and stored in a humidified incubator at 37°C, 5% CO_2_ prior to performing the ELISpot assay.

The ELISpot assay was performed using the Murine IFN-*γ* Single-Colour Enzymatic ELISPOT Assay kit (CTL). Briefly, PVDF membranes were coated with IFN-*γ* capture antibody overnight and then washed with PBS. 100 μL of test peptide (at a final concentration of 1 μg/ml) PepMixTM SARS-CoV-2 S (JPT, PM-WCPV-S-1) or murine anti-CD3 antibody (positive control) (Invitrogen, 16-0031-82) or CTL-TestTM medium (negative control) were applied to the wells and plates were then placed in a 37°C humidified 5% CO_2_. Splenocytes were plated at 4 x 10^5^ cells/well in 100 μL suspension in CTL-TestTM medium and immediately transferred to a humidified incubator at 37°C, 5% CO_2_ for 24 hours. Following development of Spot Forming Units (SFU) and prior to analysis, plates were dried overnight. The plates were scanned and analysed using the ImmunoSpot^®^ S6 Ultra-V analyzer (CTL). For all wells, the number of SFU was determined using the SmartCountTM and Autogate and expressed as SFU/million cells. Both peptide stimulation and medium treatment was carried out in triplicate for each sample.

### SARS-CoV-2 Surrogate Virus Neutralization Test

Blocking of the RBD-ACE2 interaction by the mouse sera was assessed using a SARS-CoV-2 Surrogate Virus Neutralization Test Kit (Genscript) (Tan et al. 2020), following manufacturer’s instructions. Briefly, sera from the 3rd bleed were diluted in Phosphate Buffered Saline (PBS), before further diluting in the provided sample buffer in a 1:9 ratio. Samples and controls were then mixed with HRP-RBD and incubated in 37°C for 30 minutes, and added to the ACE-2 coated wells. The plate was incubated in 37°C for 15 minutes and washed four times. TMB solution was added to the reactions, followed by a 15-minute incubation in the dark at room temperature. Absorbance at 450 nm was read immediately following quenching with the provided stop solution (Synergy H1 Hybrid Multi-Mode Reader, BioTek). The IC_50_ was estimated using a 3 parameter log-logistic regression fit in Graphpad Prism. Absorbance at 450nm was modelled in response to sera dilution.

### Peptide epitope scanning

Peptide epitope scanning was performed against a microarray of 15mer peptides with 13mer overlap covering the SARS-CoV-2 spike protein as well as other coronavirus spike proteins using the PEPperCHIP^®^ Pan-Corona Spike Protein Microarray (Dahlke et al. 2020; Heiss et al. 2020). Experiments were performed by PEPperPRINT GmbH (Heidelberg, Germany). Pre-staining of the peptide microarray was done with the secondary antibody to identify any background interactions with the 4,564 different spike protein peptides of the microarray that could interfere with the main assays. Subsequent incubation of other microarrays with mouse sera at 1:100 dilution was followed by staining with secondary and control antibodies. Read-out was performed with a LI-COR Odyssey Imaging System at scanning intensities of 7/7 (red/green). Additional HA control peptides framing the peptide microarrays were subsequently stained as internal quality control to confirm the assay performance and the peptide microarray integrity. Quantification of spot intensities and peptide annotation were performed with PepSlide^®^ Analyzer (PEPperPRINT GmbH). In Figure 7, spot intensities were plotted on a log(10) scale.

### Multiplex particle-based flow cytometry (Luminex)

Luminex assays were performed essentially as previously described in Xiong et al. 2020. RBD, S-GSAS/PP, S-R/PP (two independent preparations), S-R, S-R/x2, as well as S-R and S-R/x2 pre-incubated at 37°C for 30 minutes or 60°C for 30 minutes, were covalently coupled to distinctive carboxylated bead sets (Luminex; Netherlands) to form a 10-plex assay.

The S-variant and RBD coupled bead sets were incubated with sera from immunized mice at 4 dilutions (1:100; 1:1000; 1:10000; 1:100000) for 1 h in 96-well filter plates (MultiScreenHTS; Millipore) and analyzed on a Luminex analyzer (Luminex / R&D Systems) using Exponent Software V31. Specific binding was reported as mean fluorescence intensities (MFI). MFI values at each of the four dilutions were used to estimate IC_50_ values as described below.

### Statistical tests

Pseudovirus neutralisation and antigen binding potency were estimated using a 4 parameter log-logistic dose response curve. Dose response curves were fit using the `drc` (Ritz et al. 2015) package in R (R Core Team 2018). A single model was fit to data from all mice for a given experiment but a separate IC_50_ value was estimated for each mouse. This meant that a separate curve was fit to each mouse but they only differed in their IC_50_.

To test if S-R/x2 sera neutralised pseudovirus better than expected given its S-R/PP binding or surrogate virus neutralisation we fit linear regression models predicting pseudovirus neutralisation based on S-R/x2 status and either S-R/PP binding IC_50_ or surrogate virus neutralisation IC_50_. In these models S-R/x2 status was a binary variable indicating if the mouse was immunized with S-R/x2 or not. The significance of the S-R/x2 status was assessed by ANOVA with a simplified model were S-R/x2 status was not included. These models and comparisons were performed in R (R Core Team 2018).

## Supplementary Figures

**Figure S1.**
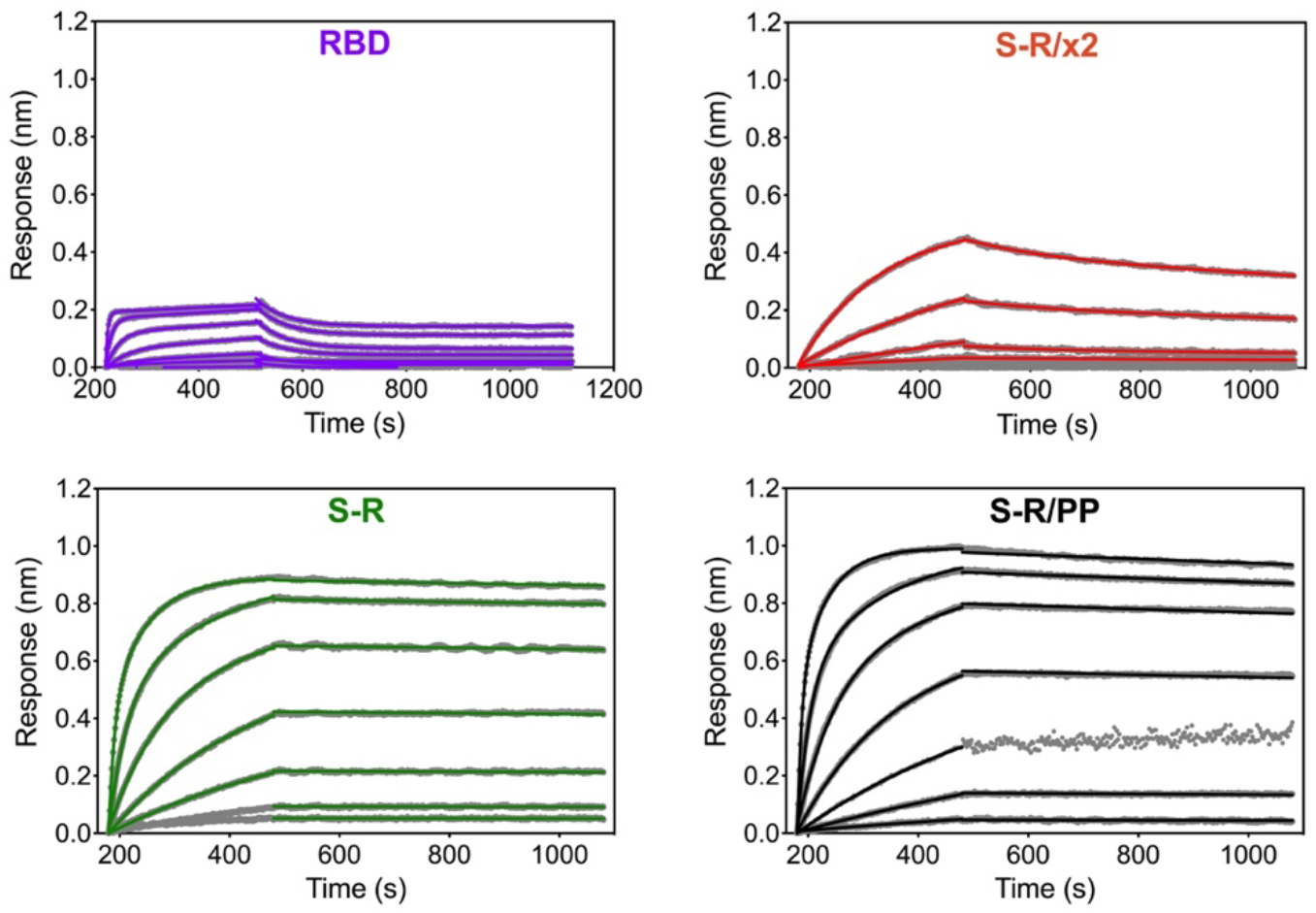
Biolayer interferometry sensograms. Biolayer interferometry sensograms for binding kinetics of ACE2 to RBD, S-R/x2, S-R and S-R/PP. Data are shown in grey with fits to the data in their respective coloured lines. The kinetic data and calculated dissociation constants (*K_d_*) are summarised in **Table 1**.

**Figure S2.**
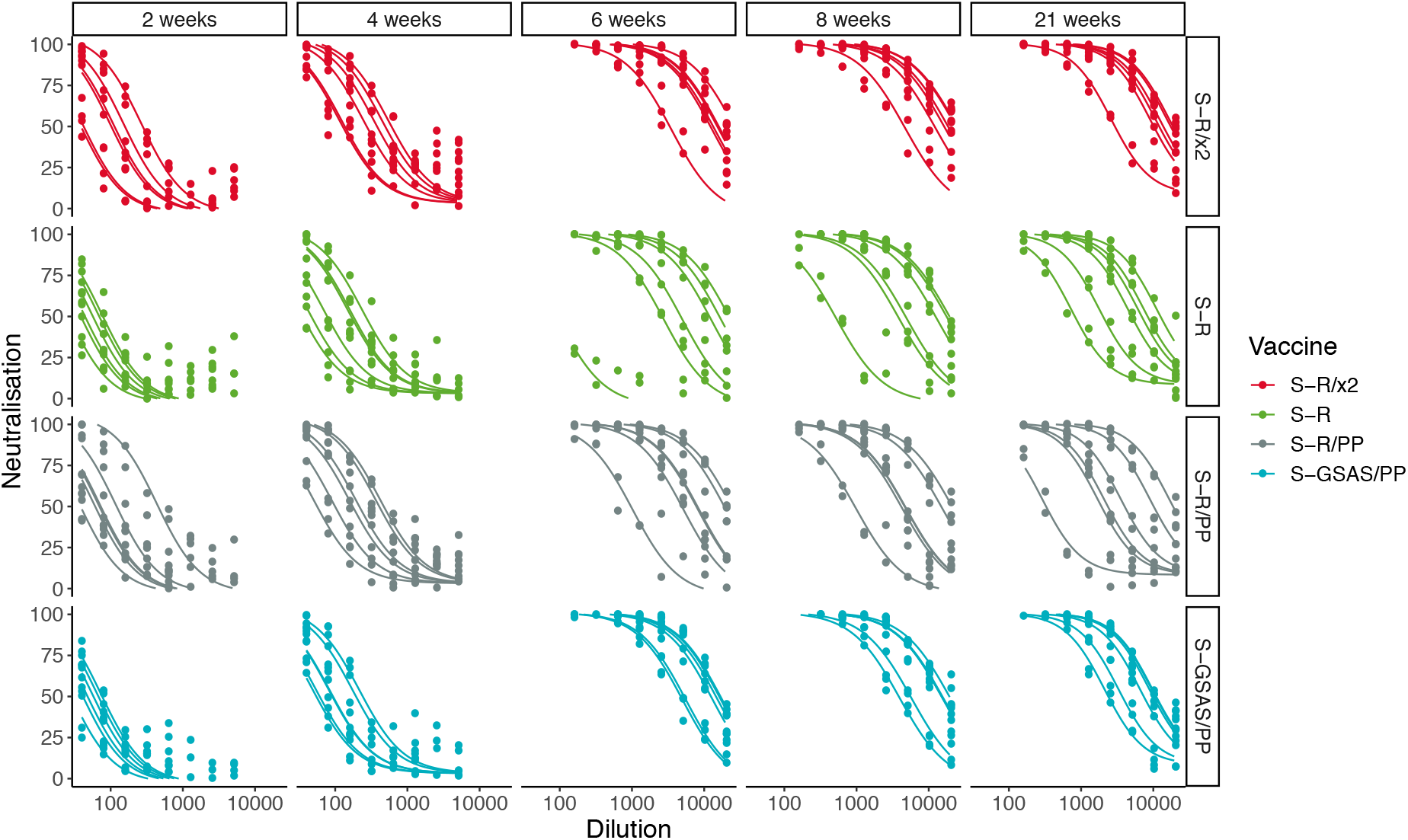
Four-parameter dose response curves for SARS-CoV-2 pseudovirus neutralisation for each individual mouse coloured by immunogen. Curves are shown for the time points, 2, 4, 6, 8, and 21 weeks post prime.

**Figure S3.**
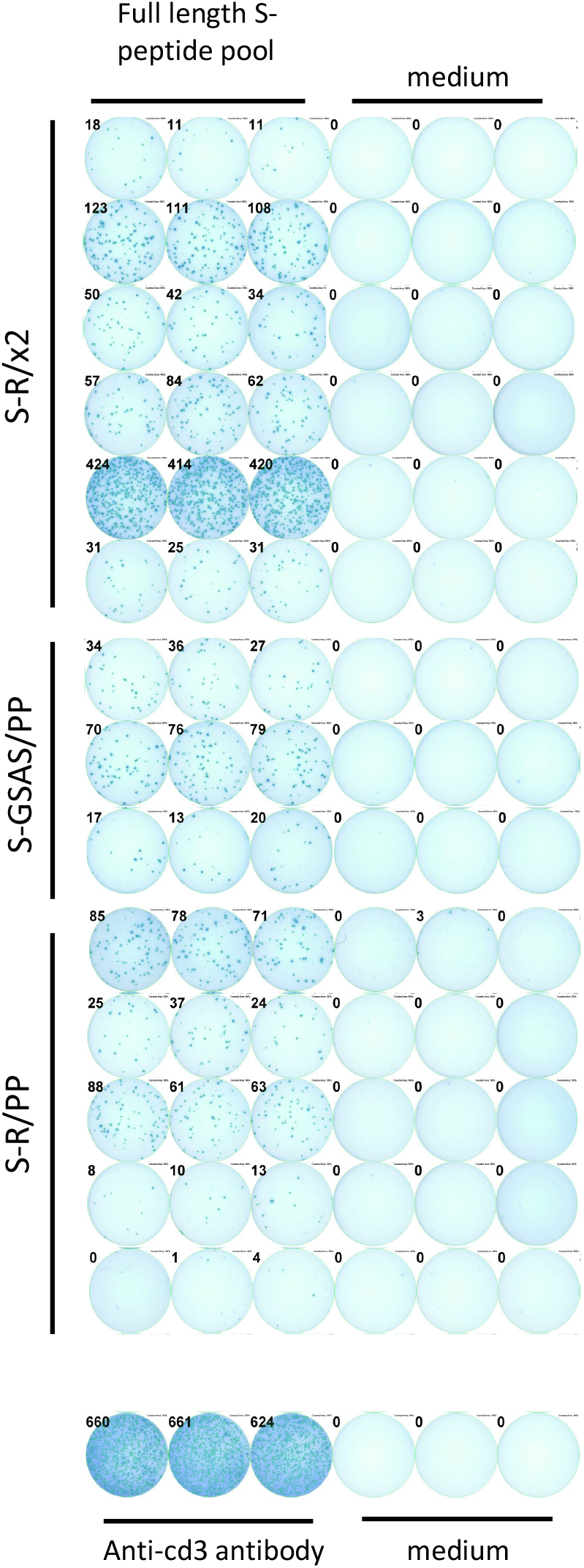
Images of IFN-*γ* mouse splenocytes ELISpot assay. Mice were immunized with the indicated constructs and SARS-CoV-2 S-specific IFN-γ secreting cells in the spleens of immunized animals were enumerated via ELISpot assay. Images show spot counts per 4 x 10^5^ splenocytes in response to S peptides, medium or anti-cd3 (positive control). Images were captured by ImmunoSpot^®^ S6 Ultra-V analyzer (CTL).

